# A stretchable and biomimetic polyurethane membrane for lung alveolar *in vitro* modelling

**DOI:** 10.1101/2024.12.09.625423

**Authors:** Emmanouela Mitta, Andrew P. Gilmore, Angeliki Malliri, Sarah H. Cartmell

## Abstract

The lung alveolus constitutes a morphologically and mechanistically complex tissue that is constantly subjected to cyclic tension and exhibits unique elastic properties. Available materials used to mimic alveolar tissue often lack biomimicry and the mechanical properties required for cyclic tension. Here, we report a fully synthetic fibrous polyurethane scaffold that approximates tissue stiffness, is elastic under breathing simulations and supports long-term culture of alveolar epithelial-like cells. Using electrospinning a fibrous membrane of tuneable thickness, set fibre diameter and small pore size is prepared. When subjected to cyclic uniaxial tension the material retains its elasticity at both low and high frequency mimicking human and mouse breathing. Thanks to the small pore size, lung alveolar cells can be cultured on its apical surface forming an epithelial monolayer. This monolayer can be maintained long term (at least 15 days) and in an air-liquid interface. In the latter conditions, cells differentiate and exhibit expression of surfactant protein A, a constituent of the surfactant layer that plays a key role in lung physiology. Owing to its lung-mimicking characteristics, the electrospun membrane holds the potential to be adapted for breathing lung models.

## INTRODUCTION

Being a respiration organ, the lung undergoes cyclic mechanical tension during breathing in order to facilitate gas exchange of O_2_ and CO_2_ in the blood. This process takes place at the alveolus, the functional unit of the organ, and is responsible for maintaining lung cell homeostasis. Abnormal mechanical loading or its absence is involved in alveolar cell population imbalance and reduced surfactant protein expression that ultimately constricts alveoli^1,2^.

*In vitro* models are a powerful tool for studying human diseases and if used as preclinical models they can minimise the use of animals in research. To achieve this and produce meaningful data that can be translated to the clinic, models are required to recapitulate the tissue or organ in question. The effect of breathing is gradually incorporated in lung research and its significance is demonstrated in lung *in vitro* models. Cell-stretching bioreactors such as organ-on-chip technology has demonstrated how cyclic stretch can reduce cancer cell growth and recapitulate the cancer growth timeframe and treatment response observed in animal models^3^. Some evidence also suggests that cyclic stretch is responsible for lowering efficacy of chemotherapy agents on cancer cells^4^, while the opposite can be observed with inhaled or orally administered drugs against idiopathic pulmonary fibrosis^5^. These models achieve organ-level architecture including air-liquid interface (ALI) and can mimic different types of cyclic stretch such uniaxial and multiaxial^6–8^. However, prototypes of these models rely on materials that lack biomimicry. One example is polydimethylsiloxane (PDMS), a flexible silicone-based material^9^ that can approximate the alveolar barrier thickness^10^ but lacks the fibrous topography of the lung extracellular matrix (ECM)^11^. It further requires protein coatings as its strong hydrophobicity hinders cell attachment^12^. As an effort to replace PDMS, natural ECM components such as collagen, gelatin and elastin have been considered as they offer the best lung ECM mimicry^13,14^. However, the scaffolds’ mechanics remain unknown, and they are met with batch variability and lack of standardisation.

Use of synthetic materials remains a promising avenue for approximating the lung mechanics. Polyurethane elastomers, a class of materials with high elasticity and vast applications including the medical field^15^, would make a suitable candidate for lung modelling. Domansky and colleagues^16^ compared the elastic properties of PDMS to those of polyurethane under human breathing conditions and found no significant differences between the two concluding that polyurethane is a desirable alternative to PDMS. According to authors, this replacement would make lung-on-chip models suitable for pharmacological applications, as contrary to PDMS, polyurethane does not adsorb small hydrophobic molecules. Polyurethane has since been explored by other groups with the aim to fabricate flexible scaffolds for lung and vocal fold tissue engineering^17,18^. In the case of lung, polyurethane has been evaluated as a 2D membrane still lacking the fibrous nature of the ECM.

In this work, electrospinning of medical-grade thermoplastic polyurethane (TPU) is employed as a method to fabricate a fibrous and porous lung-mimicking membrane. The scaffold’s elasticity is assessed by means of uniaxial tension including breathing simulation at different frequencies to test its suitability for cyclic stretch applications. Cell viability as well as epithelial monolayer development and surfactant protein expression were evaluated demonstrating TPU as a suitable lung-specific biomaterial that can be adopted by cell-stretching technologies.

## MATERIALS AND METHODS

### Electrospinning

A 3% (w/v) solution of SG-80A TPU (Tecoflex®; IMCD, UK) was prepared by dissolving SG-80A pellets in a chloroform:methanol (1:1 v/v) solvent system and stirring overnight at room temperature. The solution was electrospun using a vertical stationary spinneret setup equipped with a 23G needle and flat collector. Process parameters were set at 0.5 ml/h flow rate, 15 kV high voltage and 20 cm needle-collector distance. All electrospinning was carried out at 18-20 °C and 40-50% relative humidity.

### Morphological characterisation of the electrospun mesh

#### Fibre diameter and mesh pore size

Samples were mounted on aluminium pin stubs using conductive double-sided adhesive carbon tabs and sputter coated with a thin 3 nm layer of Au/Pd prior to imaging. Images were captured on a Vega TC (Tescan, Czech Republic) scanning electron microscope (SEM) using 200 Hz unidirectional scan speed, 1024×1024 formation. Fibre diameters were analysed using ImageJ2 (v. 2.3.0/1.53f) and their distribution was determined from a minimum of 300 fibres per replicate.

Pore size was calculated based on previous work^19^. SEM images used to measure fibre diameter were segmented and pore area was recorded in ImageJ2 (v. 2.3.0/1.53f).

#### Water contact angle

A 3% TPU solution was electrospun as described previously and allowed to dry for 24 hours. Strips of electrospun material were attached to glass slides and mounted on a contact angle microscope (KRUSS, Germany). Using a needle, 1.3 μl of distilled deionised water was carefully placed on the surface of the electrospun mesh. The contact angle on either side of the droplet were measured using the Drop Shape Analysis software and the tangent method 1.

### Mechanical characterisation of the electrospun mesh

Samples were prepared by cutting thin strips of electrospun TPU (width: 5 mm, length 25 mm) which were mounted on paper boards (width: 12.5 mm, length: 45 mm) using double-sided tape. Mounted samples were gripped between two tensile grips and sides of paper boards were cut to release the scaffold and the tension created by the board. Tensile tests were performed on an Electroforce 3310 instrument (TA instruments, USA) equipped with a 22N load cell and a strain rate of 5 mm/min using a ramp waveform setup. Electrospun meshes were kept dry or submerged in water during the tests. Stress-strain curves were plotted in Prism (Version 10, GraphPad, UK) and the Young’s modulus was determined by calculating the slope of the stress-strain curve.

To assess elasticity during physiological breathing conditions, cyclic tensile tests were carried out on a Planar Biaxial instrument (TA Instruments, USA) equipped with a 5N load cell. A sinusoidal wave was applied to mimic breathing motion (10% strain, 0.2 Hz for human breathing; 10% strain, 1.3 Hz for mouse breathing^20^), which lasted 1 hour. All tests were carried out under dry conditions at room temperature.

### Cell line and culture

#### Cell line and maintenance

The human lung cancer cell line NCI-H441 (H441) purchased from ATCC was cultured in RPMI-1640 (A1049101, Thermo Fisher Scientific, UK) supplemented with 10% Fetal Bovine Serum (FBS FB-1345/500-S00Q2, LabTech). Cells were maintained at 37°C, 95% air and 5% CO_2_. For surfactant protein expression monitoring, H441 cells were cultured in liquid-liquid interface (LLI) for 4 days to reach confluency and then transferred to ALI for 6 days by removing media from the apical compartment of cell culture inserts.

#### TPU attachment and preparation for cell culture

To assess biocompatibility of the electrospun TPU scaffold, membranes were mounted on cell culture inserts. Following electrospinning on baking paper, the membrane was stored in a fume hood for at least 24 hours for solvents to evaporate. The membrane was then mounted using silicone adhesive (Corning 732, Corning, USA) on glass coverslips or Thincert^TM^ inserts (#662641, Greiner Bio-One, UK) after cutting out the PET membrane supplied (Figure S1A). Coverslips and inserts were left to dry overnight at room temperature and were then sterilised under ultraviolet (UV) light for 20 minutes on either side. Finally, TPU scaffolds were washed in 5% Gentamicin solution (G1272, Sigma-Aldrich, Germany) followed by one wash in phosphate buffered saline (PBS; D8537, Sigma-Aldrich, Germany) and incubated with RPMI-1640 (supplemented with 10% FBS) at 37° C overnight before seeding (Figure S1B). Thincert^TM^ inserts with a 0.4 μm pore PET membrane and glass coverslips were used as control where appropriate.

#### Seeding density optimisation

For seeding optimisation, H441 cells were stained in solution prior to seeding using the Vybrant™ DiD cell-labelling solution (V22887, Invitrogen, Germany) according to manufacturer’s instructions. H441 cells were seeded on polyethylene terephthalate (PET) or TPU cell culture inserts at 2.5×10^4^, 5×10^4^ or 10^5^ cells/cm^2^ and growth was monitored over 11 days. Images were captured on an EVOS™ M7000 Imaging System (Thermo Fisher Scientific, UK).

### Cell viability

#### Viability/Cytotoxicity

To visually assess cell viability and cytotoxicity on the electrospun TPU following seeding, the LIVE/DEAD™ viability/cytotoxicity kit (L3224, Thermo Fisher Scientific, UK) was used. H441 cells were seeded on uncoated or TPU-coated glass coverslips at 5×10^4^ cells/cm^2^ and were allowed to adhere overnight. The following day, media was aspirated, and cells were washed once in PBS. LIVE/DEAD™ working solution (2 μM calcein AM and 4 μΜ EthD-1 in PBS) was added and cells were incubated for 20 minutes at 37 °C. LIVE/DEAD solution was then removed, and samples were washed once in PBS before mounting on glass slides for imaging. Images were captured on the EVOS™ M7000 Imaging System (Thermo Fisher Scientific, UK).

#### Cell metabolism assay

Cell viability was monitored over time by measuring cell metabolic activity. H441 cells were seeded in 24-well cell culture inserts with a PET or TPU membranes at a density of 5×10^4^ cells/cm^2^. A 10X alamarBlue^TM^ solution (BioRad, UK) was diluted 1:10 in RPMI-1640 cell culture media. 200 μl of the solution was added to each insert and samples were incubated at 37° C for 2 hours. Upon completion of the incubation, 100 μl of the supernatant was transferred to a 96-well plate. Inserts were washed once in PBS and media were replenished until the next timepoint. Fluorescence was measured at 582 nm on a FLUOstar Omega microplate reader (BMG Labtech). Background subtraction was achieved using blank values from PET or TPU inserts without cells.

#### Double-stranded DNA quantification

Cell proliferation was evaluated by measuring the double-stranded (ds) DNA content on TPU versus PET scaffolds. H441 cells were seeded in 24-well cell culture inserts with a PET or TPU substrate at a density of 5×10^4^ cells/cm^2^. The dsDNA was isolated and quantified using the Quant-iT™ PicoGreen™ dsDNA assay kit (P7589, Thermo Fisher Scientific, UK) following the manufacturer’s protocol. Fluorescent signal (Excitation: 488 nm; Emission: 520 nm) was measured on a FLUOstar Omega microplate reader (BMG Labtech).

### Epithelial monolayer development

#### Cell Fixation

Cells seeded either on PET (5×10^4^ cells/cm^2^) or TPU inserts (10^5^ cells/cm^2^) were washed with 1X PBS and fixed in 4% Paraformaldehyde (#28908, Thermo Fisher Scientific, UK) for 10 minutes. Cells on TPU inserts were fixed for 20 minutes. After fixation, samples were washed three times in PBS and permeabilised with 3% bovine serum albumin (A7906, Sigma Aldrich, Germany). Samples were stored in PBS at 4° C until further analysis.

#### Immunofluorescent staining

Upon staining, samples were blocked in antibody dilution buffer (ADB; 0.02% Triton X-100, 5% horse serum) for 10 minutes in a dark humidity chamber followed by incubation with primary antibodies ZO-1 (1:150, #40-2200, Invitrogen, Germany) and E-cadherin (1:400, #610181, BD Transduction Labs, USA) diluted in ADB for 2 hours at room temperature. Samples were then washed three times in PBS and further incubated with secondary antibodies Alexa Fluor 488 (1:500, A11029, Invitrogen, Germany) and Alexa Fluor 594 (1:500, A11037, Invitrogen, Germany) that were also diluted in ADB for 1h at room temperature in the dark. Upon completion, samples were washed three times in PBS and incubated with 0.1 μg/ml DAPI for 10 minutes and rinsed in distilled deionised water. Finally, coverslips and membranes were left to dry overnight. TPU and PET membranes were then cut carefully using forceps and were mounted on glass slides using DAKO Mounting Medium and square glass coverslips in a sandwich. Cell-laden coverslips were mounted cell-side down using the same mounting media. Slides were dried at room temperature overnight before imaging.

#### Fluorescent imaging

Images were acquired on a Leica TCS SP8 upright confocal microscope (Leica, UK) using 63x/1.40 oil immersion lens. Images were captured with the following settings: pinhole 1 airy unit, scan speed 200 Hz unidirectional, formation 1024×1024. On the EVOS, 10x, 20x LWD and 40x LWD lenses were used. Acquired images were processed in ImageJ Fiji (Version 2.3.0).

#### RNA extraction and cDNA synthesis

To measure mRNA expression of SP-A, RNA was extracted from H441 monolayers in ALI. Cell monolayers were washed twice in RNase-free PBS and total RNA was extracted using the Qiashredders (#79656, Qiagen, UK) and RNeasy mini kit (#74104, Qiagen, UK) following the manufacturer’s protocol. RNA concentration and quality were determined using a Nanodrop 2000 (Thermo Fisher Scientific, UK) and samples were stored at −80° C until further analysis. For quantitative polymerase chain reaction (qPCR) experiments, cDNA was synthesised using the High-Capacity RNA-to-cDNA™ kit (#4387406, Applied Biosystems, USA).

#### qPCR

qPCR was performed using the Luna® universal qPCR master mix (M3003, New England Biolabs, UK) following the manufacturer’s protocol on a StepOnePlus qPCR instrument (Applied Biosystems, USA). Primers were designed against the *SFTPA1* and housekeeping *GAPDH* genes (Table 1) using the primer-BLAST primer designing tool. Primer sequences and melting temperatures (Tm) can be found in Table 1. Thermal cycling conditions were set according to manufacturer’s instruction and are summarised in Table 2. Mean C_T_ values were extracted and converted to ΔΔC_T_ values which were further converted to 2^-ΔΔCT^ and expressed relative to the 2^-ΔΔCT^ of the LLI condition.

**Table 1.**
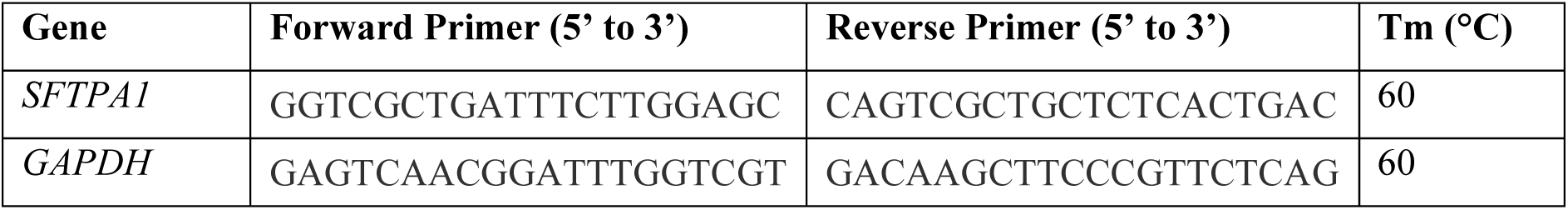
qPCR primers.

**Table 2.** qPCR thermal cycling conditions.

| Cycle Step | Temperature (°C) | Time (s) | Cycles |
| --- | --- | --- | --- |
| Initial Denaturation | 95 | 60 | 1 |
| Denaturation | 95 | 15 | 40 |
| Anneal | Variable (See $T_m$) | 30 | |
| Extend | 72 | 60 |  |

#### Transepithelial Electrical Resistance measurements

To assess the quality of the epithelial barrier, the transepithelial electrical resistance (TEER) was measured on confluent cell layers. H441 cells were seeded at 5×10^4^ or 10^5^ cells/cm^2^ in 24-well cell culture inserts with a PET or TPU substrate. Prior to TEER measurements, media in well and insert were replenished, and cells were returned to the incubator (37° C, 5% CO_2_) for 15 minutes. TEER was measured using an EVOM2 epithelial Volt/Ohm meter (World Precision Instruments, USA) equipped with an STX3 pair of electrodes. Electrodes were submerged in 70% ethanol and PBS prior to the session and in between measurements of different samples. Two different readings were collected from two edges of each insert and their average was calculated. All TEER measurements were performed with samples at 35-37° C and each round of measurements lasted up to 5 minutes. Cells were returned to the incubator for 5 minutes and retrieved for subsequent rounds of measurements.

### Statistical analysis

Statistical analysis was carried out in GraphPad Prism (Version 9.4.0). Bar and XY plots show mean, and error bars indicate the standard deviation (SD). Statistical comparison was performed using two-tailed unpaired t test with Welch’s correction for comparisons of two independent groups, or one-way analysis of variance (ANOVA) with Tukey’s post-hoc test for comparisons of three or more independent groups, or two-way ANOVA with Holm-Šidak post-hoc test for comparisons of two or more independent groups with two independent variables. Statistical tests and their results are detailed in the figure legends. Differences between groups were considered statistically significant when p<0.05. Not significant (ns) differences are indicated in the graphs.

### Data availability

The datasets generated during and/or analysed during the current study are available from the corresponding author on reasonable request.

## RESULTS

### Electrospun TPU scaffold characterisation

Upon electrospinning of TPU and collecting on a static surface, fibrous scaffolds with randomly aligned nanofibers can be obtained (Figure 1A), with tuneable thickness. For the purpose of experiments detailed in this paper, 10 μm-thick membranes were fabricated (Figure 1B) with an average fibre diameter of 0.44±0.14 μm (Figure 1C) and average pore size 1.03±0.29 μm (Figure 1D). To calculate the hydrophobicity of the TPU under dry conditions the mean contact angle of a water droplet placed on a TPU film was calculated by averaging the contact angles either side of the droplet. TPU was deemed hydrophobic with a mean contact angle of 116.4±17.6° (Figure 1E).

**Figure 1.**
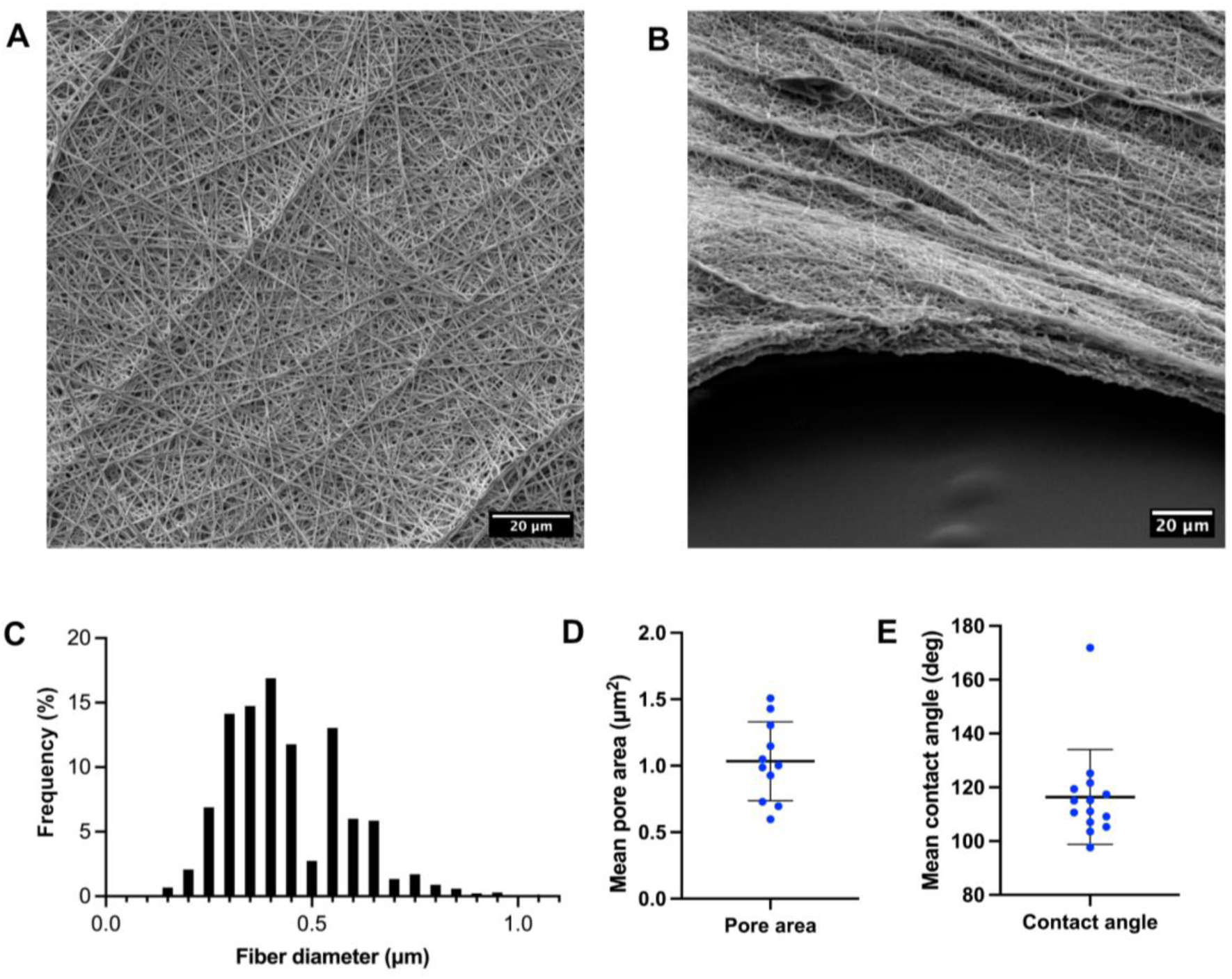
**A-B.** SEM images of electrospun TPU. **(A)** shows the randomly aligned mesh of nanofibres that can be achieved and **(B)** the three-dimensional surface topology. **C-E.** Morphological characterisation of the electrospun TPU mesh showing **(C)** fibre diameter frequency distribution of TPU fibres shown in (A). Mean fibre diameter 0.442±0.14 nm. **(D)** Pore area calculated from SEM images using a thresholding method. Mean pore size 1.035±0.9 μm^2^. **(E)** Contact angle measurements of dry electrospun TPU. Mean contact angle 116.2°±17.64 Each point represents a technical replicate, n=3. Scale bars: 20 μm.

As shown in Figure 2A, the electrospun scaffold has appropriate mechanical properties for its lung tissue application (Young’s modulus = 11.3±6.5 MPa) demonstrating a longer elastic period and a short plastic period on the stress-strain curve. Simulation of the aqueous conditions in cell culture results in a softer material with significantly lower Young’s modulus (Figure 2B) that retains a similar tensile profile. Mean Young’s modulus, ultimate stress and ultimate strain values are summarised in Table 3.

**Figure 2.**
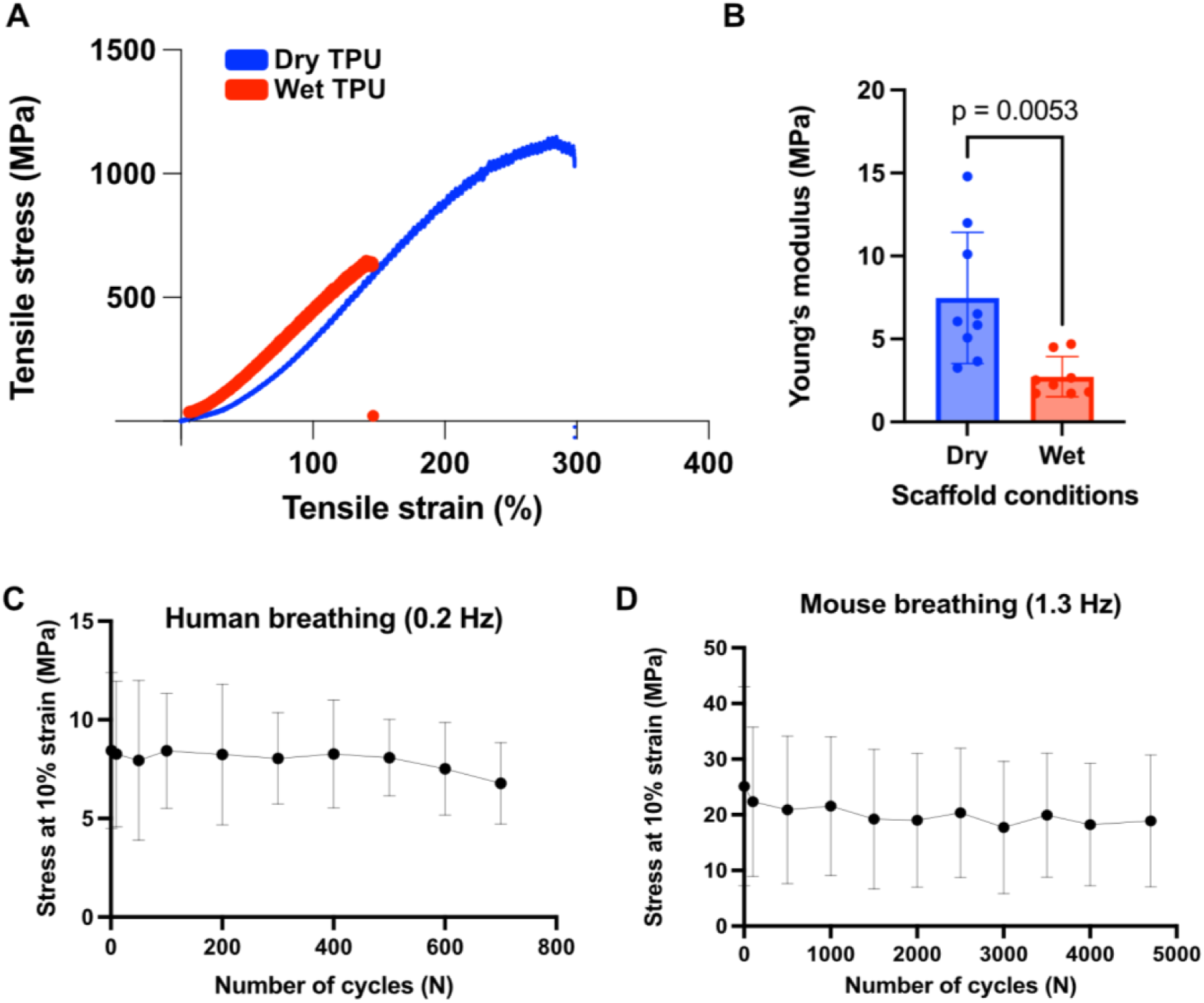
**(A)** Representative stress-strain plots of electrospun TPU subjected to uniaxial tension under dry (blue) or wet (red) conditions. For wet conditions material was submerged in aqueous solution during testing. (**B)** Young’s moduli derived from stress-strain curves comparing dry to wet scaffolds. Each point represented a technical replicate. Statistical significance was determined by an unpaired two-tailed t test, n=3. **C-D.** Elasticity of electrospun TPU plotted as the mean stress at 10% strain against the number of cycles equal to 1 hour of cyclic tension. Scaffolds were stretched at the **(C)** human (0.2 Hz) or **(D)** mouse (1.3 Hz) breathing frequency resulting in 720 and 4,700 cycles, respectively. Mean stress: 8.86±0.15 MPa (human), 20.29±2.12 MPa (mouse). Mean

**Table 3.** TPU mechanical properties (Mean ± SD)

|  | **Dry TPU** | **Wet TPU** |
| --- | --- | --- |
| **Young's modulus (MPa)** | 7.49±3.4 | 2.73±1.2 |
| **Ultimate tensile stress (MPa)** | 976±29 | 634±73 |
| **Ultimate tensile strain (%)** | 311±15% | 126±16 |

### Scaffold suitability for lung breathing applications

In order to test the electrospun TPU’s suitability for breathing simulation, the scaffold was subjected to cyclic uniaxial tension, monitoring its stress levels during human and mouse breathing simulations for one hour (Figure 2C and 2D). In both cases, between cycle 1 and 50, we observe softening of the material shown as a drop in the level of strain which remains steady for the remainder of the testing period indicating that the membrane maintains its elastic properties over time. The scaffold eventually adapts to the cyclic deformation by maintaining stable stress levels thereafter. Under human breathing conditions the average stress experienced is 8.86±0.15 MPa, which increases by almost 2.5-fold when strain frequency is increased to mimic mouse breathing conditions. Despite this, no fluctuations of stress are observed at the respective conditions indicating that the scaffold can maintain elasticity over thousands of strain cycles.

### Cell viability

To test the scaffolds suitability for an alveolar epithelial model, the H441 cell line was employed. Within a day of culturing cells on the electrospun TPU, good cell attachment can be observed with minimal cytotoxicity (Figure 3A and 3B). Compared to glass, better cell attachment can be achieved on the electrospun TPU despite its inherent hydrophobicity. To assess long-term culture, a metabolic assay was used that allows continuous monitoring of cells’ metabolic activity. This showcased that H441 cells can be cultured on the electrospun TPU long-term, up to 15 days that the assay lasted (Figure 3C). During that period, a steady state of metabolic activity can be observed in H441 cells seeded on the TPU scaffold. In contrast, cells on the PET display a peak in metabolic activity by day 7 that subsequently declines and plateaus by day 15. This decline coincides with detachment of cells from the PET scaffold that are removed during the assay’s wash steps (Figure S2A).

**Figure 3.**
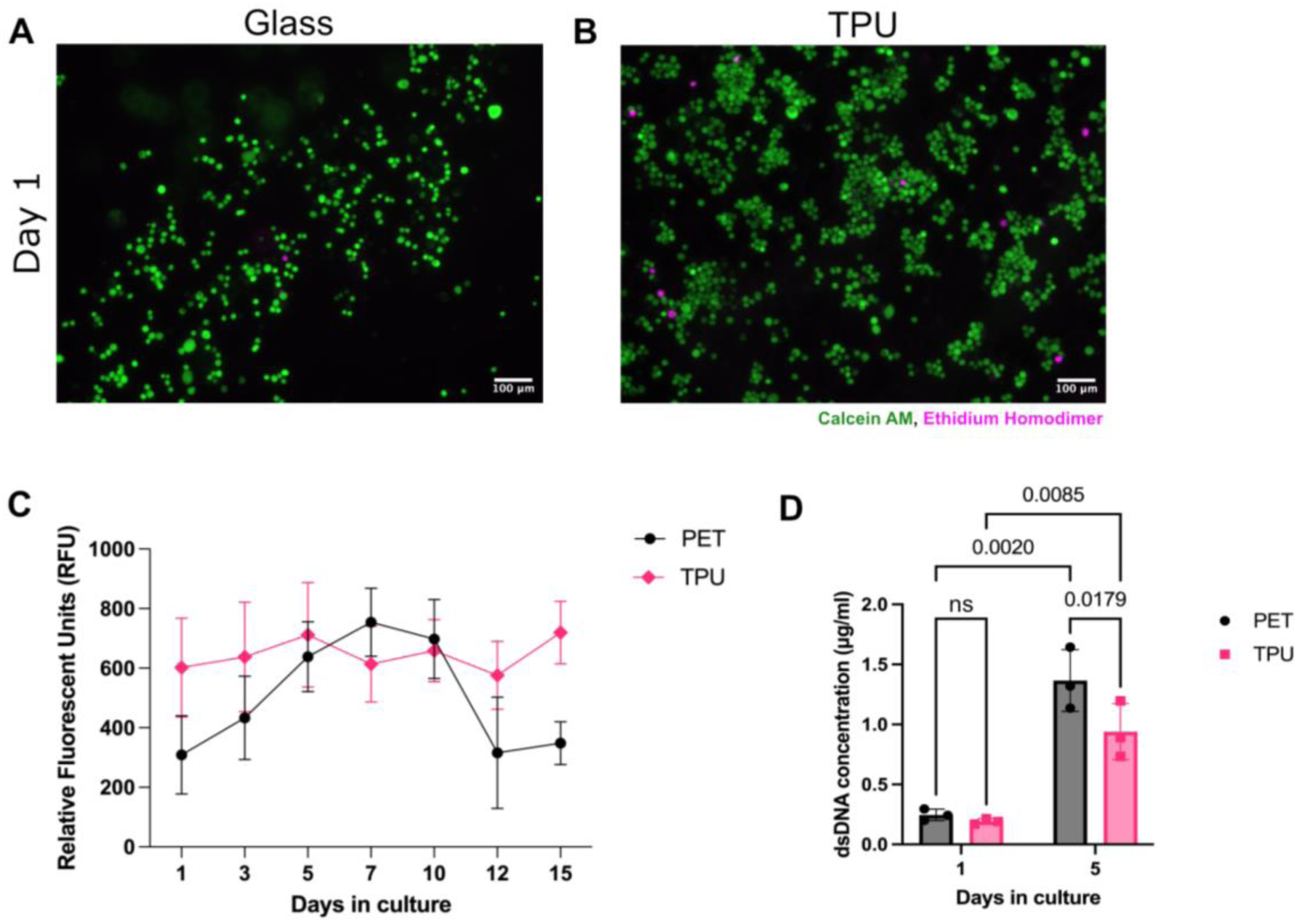
**A-B.** Representative fluorescent images of H441 cells seeded on **(A)** uncoated, or **(B)** TPU-coated glass coverslips for 24 hours and subsequently stained with the LIVE/DEAD™ viability/cytotoxicity assay kit. Live cells (green) are stained with calcein AM and dead cells (magenta) are stained with ethidium homodimer. **(C)** Cell metabolism timecourse of H441 cells seeded at 5×10^4^ cells/cm^2^ on cell culture inserts with a PET or TPU membrane. Metabolic activity was measured by means of fluorescent intensity (RFU) of the reduced alamarBlue™ reagent. **(D)** Quantification of dsDNA content of H441 cells seeded at 5×10^4^ cells/cm^2^ on PET or TPU and cultured for 1 or 5 days. dsDNA content is plotted as the mean concentration per sample per day. Statistical significance was determined by a two-way ANOVA and Holm-Šidak post-hoc test, n=3. Scale bars: 100 μm.

To confirm cell proliferation by means of metabolic activity, the dsDNA content was measured on day 1 and 5 of culture (Figure 3D). Here, no difference in dsDNA content is observed between PET and TPU on day 1 post seeding. By day 5, overall dsDNA content has increased on both membranes compared to day 1, indicative of an increased number of cells present. As expected, dsDNA content is higher on the PET membrane than the TPU showing that H441 cells on PET proliferate more than cells on the electrospun TPU scaffold.

### Epithelial monolayer development

When modelling the alveolar epithelium at least two factors should be taken into account; whether the cells can form an epithelial monolayer that mimics the alveolar barrier and whether cells can be differentiated to produce elements of the surfactant layer under ALI conditions. For this, a short differentiation protocol was adopted that includes growing H441 cells on cell culture inserts to confluency and then introducing ALI to observe surfactant production (Figure 4A).

**Figure 4.**
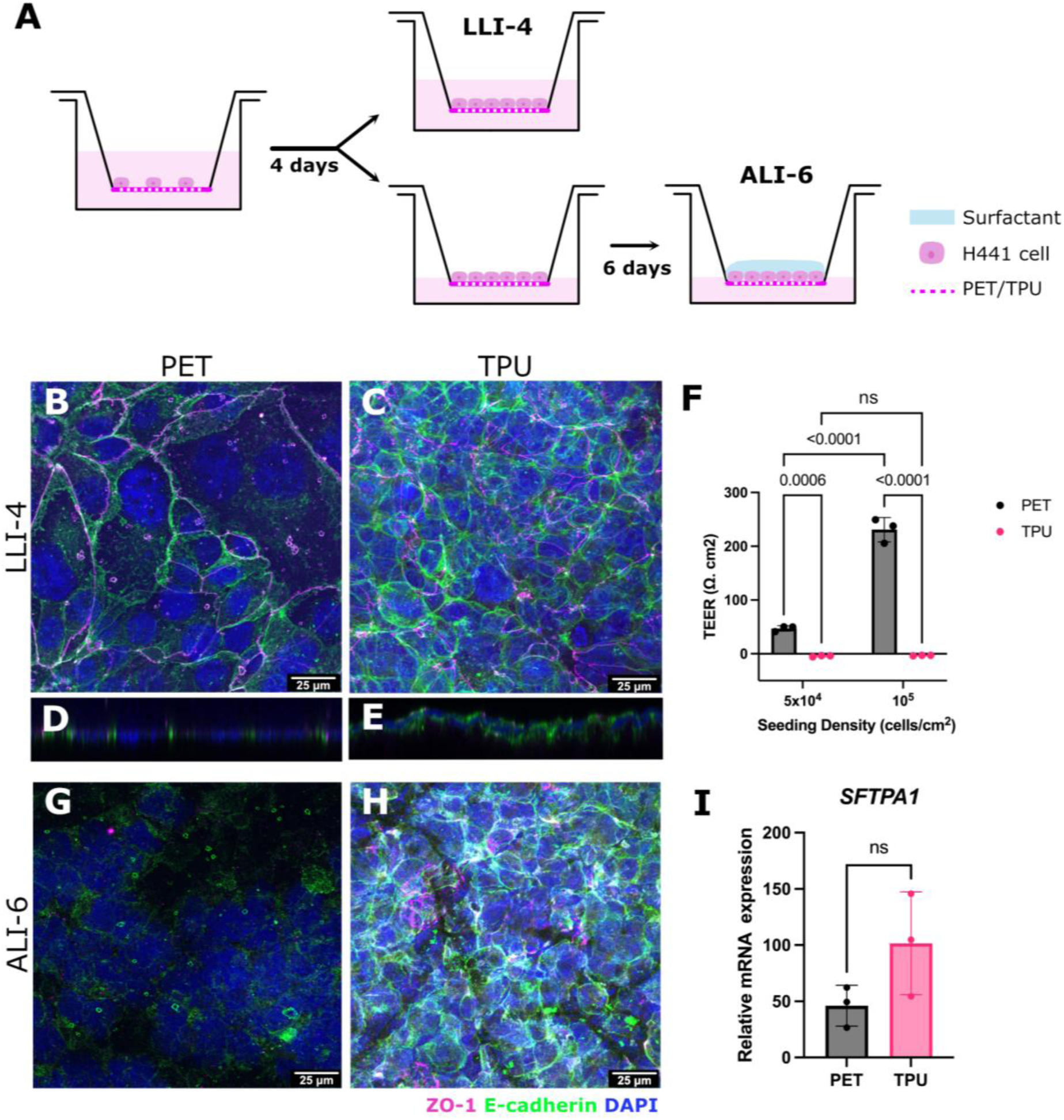
**(A)** Schematic representation of H441 differentiation in ALI conditions. H441 cells are seeded and grown to confluency within 4 days. On day 4, cells are either fixed obtaining LLI-4 samples or culture is continued in ALI. Apical removal of media leads to ALI culture and surfactant layer formation within 6 days. **B-E** and **G-H.** H441 cells immunostained for cell-cell junction markers E-cadherin (green) and ZO-1 (magenta) and nucleus (blue). H441 cells on **(B)** PET and **(C)** TPU reach confluency by day 4 and form mature cell-cell junctions. **(D-E)** Cross-sections show that H441 cells form monolayers on both substrates. **(F)** TEER measurements of H441 cells seeded on PET or TPU substrates at different seeding densities taken at LLI-4 timepoint. Upon introduction of ALI, **(G)** H441 cells on PET lose their cell-cell junctions compared to **(H)** H441 cells on the TPU substrate that maintain E-cadherin staining and ZO-1 to some extent. **(I)** mRNA expression of SFTPA1 gene shown relative to mRNA expression in LLI. No significant difference is observed between H441 cells on PET and TPU substrates by ALI6. Statistical significance was determined by two-tailed unpaired t-test, ns p>0.05, n=3. Scale bars: 25 μm.

To test epithelial monolayer formation, localisation of cell-cell junction markers E-cadherin and ZO-1 were investigated by means of immunostaining. Owning to their epithelial-like characteristics, H441 cells showcase plasma membrane localisation for both markers when seeded on PET membranes (Figure 4B). This phenotype can be faithfully replicated on the electrospun TPU scaffold within the same time frame by increasing the seeding density (Figure 4C). On PET, cells appear to cover larger surface area compared to TPU, an observation that could be attributed to differences in surface topology between the two membranes, with fibres potentially restricting cell spreading. In both cases, H441 cells form a monolayer of cells as can be seen by the cross sections taken (Figure 4D and E).

To assess the barrier properties of the H441 monolayers, TEER measurements were taken on day 4 in LLI examining two different cell densities (Figure 4F). As expected, H441 cells on the PET substrate display increased TEER that is proportional to the increase in cell density. In contrast, cells on the electrospun TPU scaffold display zero or subzero TEER values despite formation of mature cell-cell junctions. A minimal increase in TEER is observed when H441 cells are seeded at 2×10^5^ cells/cm^2^ and cells are incubated on the TPU for longer (Figure S2B).

To test SP-A expression on the electrospun TPU scaffold H441 cells were cultured in an ALI over a period of 6 days following monolayer formation. During differentiation, H441 cells on the PET lose expression of ZO-1 and E-cadherin (Figure 4G) compared to LLI culture, while cells on the TPU scaffold retain their cell-cell junctions to a good degree (Figure 4H). Cells on both control and TPU membranes showed increased expression of SP-A in response to ALI (Figure 4I) with levels similar between PET and TPU materials.

## DISCUSSION

The field of lung *in vitro* modelling has seen significant advances in the technologies used to simulate the breathing motion, but materials remain unrealistic and poorly characterised for their intended use. In this study, the electrospun TPU membrane replicates the elastin-collagen fibrous network and adheres to the tissue’s mechanical properties. Electrospinning was chosen over other fabrication methods such as spin coating as it provides a scaffold that resembles the fibrous morphology of the ECM and provides several anchorage points for cell adherence compared to a flat membrane that demonstrates a lower surface area-to-volume ratio^21^. The interconnected pore system improves wettability^22^ and allows for complete wetting of the electrospun TPU in cell culture media. The dry TPU is stiffer than the native tissue but is softened by submerging in aqueous solution. Water has been shown to soften TPU by decreasing the amount or altering the type of hydrogen bonds present within the material. It is also expected that a higher temperature (e.g. 37°C) will have an additive effect on the TPU softening when combined with water^23^.

Similarly to elastin, which can sustain up to 200% strain^23^, the electrospun TPU can achieve at least 100% elongation prior to rupture. At the physiological strain level, the elastic properties are maintained at both low and high frequencies offering the ability to model breathing patterns of different species as well as exercise. Strain frequency positively correlates with tensile stress, in line with previous findings^24^, but this does not affect the elastic properties as no reduction in mean stress is observed.

To model the alveolar tissue, the H441 cell line is employed. This is a cost-effective model of the alveolar epithelial cell that has been employed in electrospun biomaterial validation^25,26^. The fibre organisation coupled with the relatively small pore size created by the interconnected nanofibers, allows for culture of epithelial cells on the surface of the membrane, mimicking the monolayer of alveolar epithelial cells observed *in vivo*^27^. Despite the scaffold’s hydrophobicity, wetting and overnight incubation with cell culture media is sufficient to achieve cell attachment that outperforms standard commercial alternatives such as glass and PET. This comes as no surprise since fibrous topographies are shown to allow for enhanced cell adhesion compared to non-fibrous planar surfaces^28^. Although cells grow quicker on the PET membranes compared to TPU, they tend to become overconfluent and inevitably detach from the substrate. This growth discrepancy can be overcome by increasing the cell seeding density on the TPU membrane, so that H441 cells are able to form a confluent cell monolayer on the TPU which can be maintained for a longer period including in ALI. Surfactant production is key in lung physiology as it eases surface tension and prevents alveoli from collapsing^29^. In the latter conditions, it is demonstrated that a similar expression of SP-A can be achieved between PET and electrospun TPU further reinforcing the suitability of the electrospun TPU for lung cell differentiation.

The material holds the potential to culture endothelial cells in a similar fashion on the basolateral surface thus creating a model of the gas-blood barrier. It is believed that this will reinforce the barrier formation and increase TEER values currently weak in the TPU compared to PET.

In conclusion, this study explores the use of TPU as a suitable lung-mimicking material to replace current standard of practice alternatives that lack the mechanical and structural complexity of the lung ECM. Being an elastomer, TPU is an excellent candidate for lung tissue engineering and coupled with electrospinning can offer a suitable alternative for lung ECM modelling that can be adapted in existing scaffold stretching technologies including recent pressure-based approaches^25^.

## Acknowledgments

Authors would like to thank Professor Nigel Hooper and the Hooper Lab for accessing and training on the TEER Volt/Ohm meter.

## Supplementary Material

**Figure S1.**
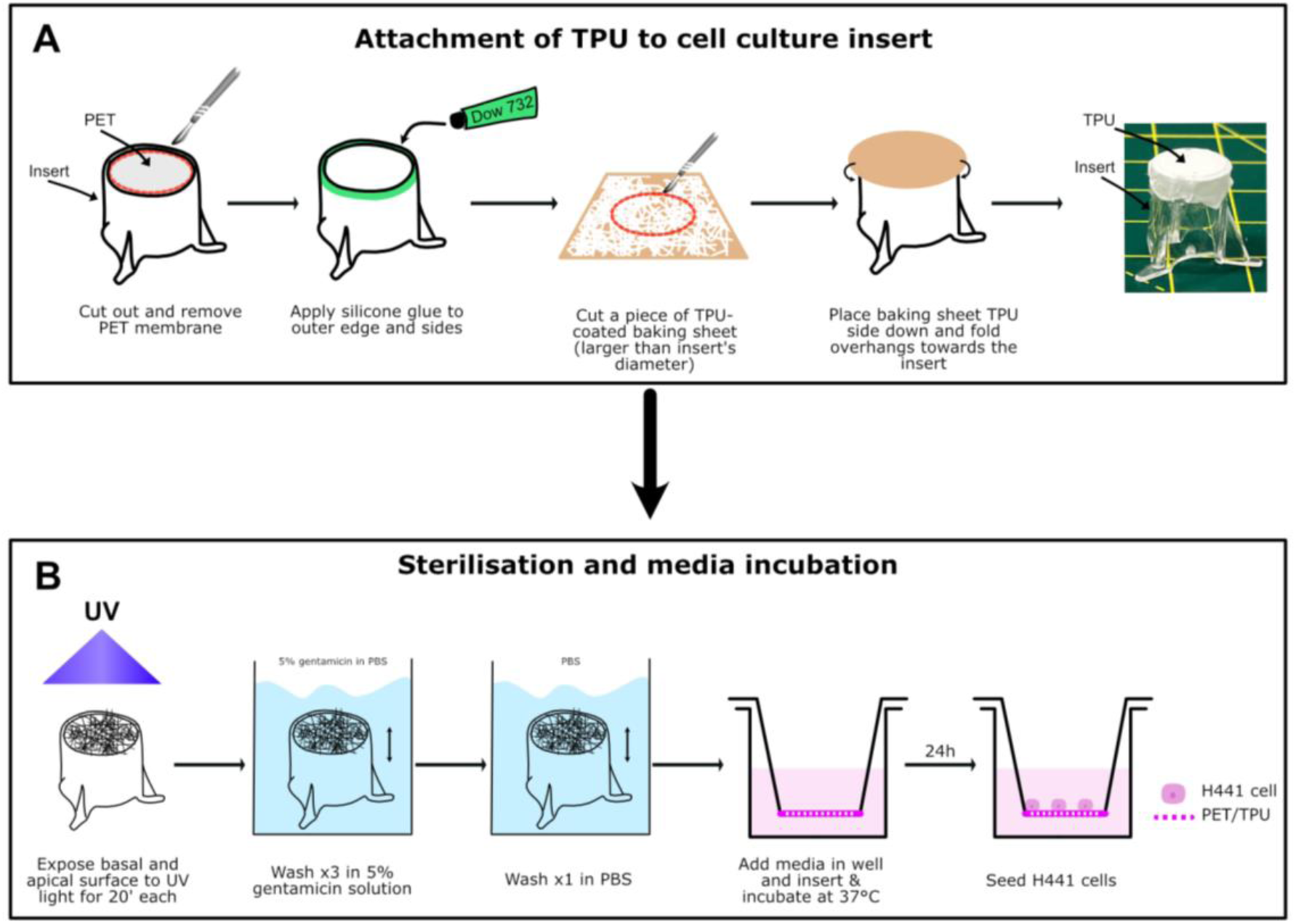
**(A)** Process of attachment of electrospun TPU on cell culture inserts. Supplied PET membrane is removed with a scalpel and silicone adhesive is applied on the outer edges of the insert. A suitable round piece of TPU-coated baking sheet is cut out and placed TPU side down on top of the insert. Overhanging sides are folded down towards the insert. Carefully the baking sheet is removed leaving the TPU only attached to the insert. **(B)** Process of TPU sterilisation once on the insert. TPU inserts are exposed to UV light for 20 minutes either side (apical and basal). Then they are transferred and submerged to a 5% gentamicin solution three times to wash. Gentamicin is washed off in PBS once and inserts are transferred to appropriate plates and incubated with media for 24h. Once incubation is complete, scaffolds are seeded with the H441 cell line at appropriate seeding densities.

**Figure S2.**
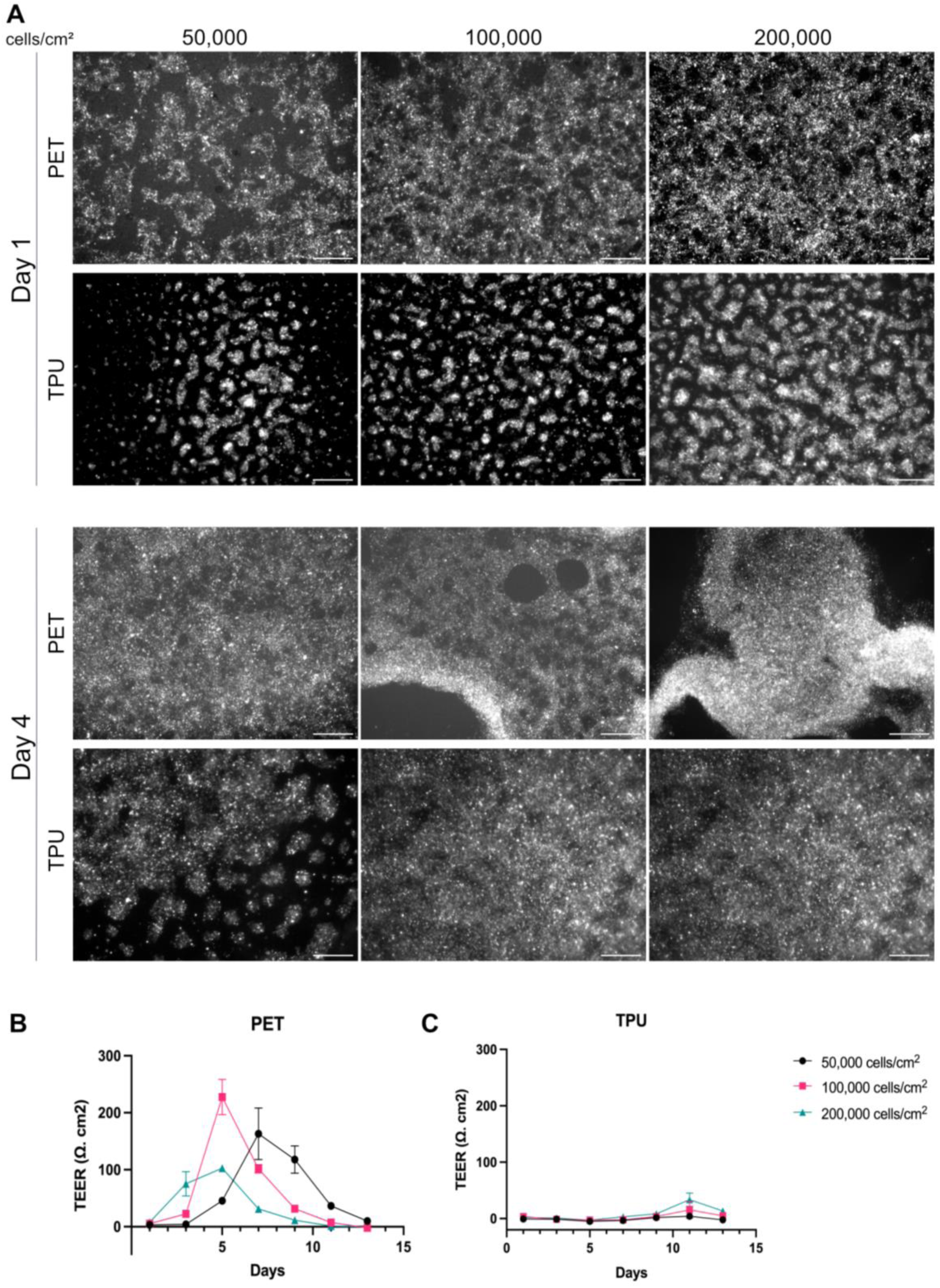
**(A)** Representative fluorescent images of H441 cells seeded at different seeding densities on PET and TPU membranes on day 0 comparing day 1 to day 4 in culture. **B-C** Monitoring of barrier integrity of H441 cells seeded at different seeding densities on **(B)** PET, or **(C)** TPU scaffolds monitored over a period of 13 days (Mean ± SD). Scale bars: 500 μm.

